# Bioimpedance and new onset heart failure: A longitudinal study of ~500,000 individuals from the general population

**DOI:** 10.1101/183483

**Authors:** Daniel Lindholm, Eri Fukaya, Nicholas J. Leeper, Erik Ingelsson

**Author notes:** Equal contribution (Nicholas J. Leeper, Erik Ingelsson). **Corresponding author:** Erik Ingelsson, Dept. of Medicine, Division of Cardiovascular Medicine, Stanford University School of Medicine, 300 Pasteur Dr, mail code: 5773, Stanford, CA 94305, USA.

## Abstract

**Importance:** Heart failure constitutes a high burden on patients and society, but although lifetime risk is high, it is difficult to predict without costly or invasive testing. Knowledge about novel risk factors could enable early diagnosis and possibly preemptive treatment.

**Objective:** To establish new risk factors for heart failure.

**Design:** We applied supervised machine learning in UK Biobank in an agnostic search of risk factors for heart failure. Novel predictors were then subjected to several in-depth analyses, including multivariable Cox models of incident heart failure, and assessment of discrimination and calibration.

**Setting:** Population-based cohort study.

**Participants:** 500,451 individuals who volunteered to participate in the UK Biobank cohort study, excluding those with prevalent heart failure.

**Exposure:** 3646 variables reflecting different aspects of lifestyle, health and disease-related factors.

**Main Outcome:** Incident heart failure hospitalization.

**Results:** Machine learning confirmed many known and putative risk factors for heart failure, and identified several novel candidates. Mean reticulocyte volume appeared as one novel factor, and leg bioimpedance another; the latter appearing as the most important new factor. Leg bioimpedance was significantly lower in those who developed heart failure (p=1.1x10^-72^) during up to 9.8-year follow-up. When adjusting for known heart failure risk factors, leg bioimpedance was inversely related to heart failure (hazard ratio [95%CI], 0.60 [0.48–0.73]) and 0.75 [0.59–0.94], in age- and sex-adjusted and fully adjusted models, respectively, comparing the upper vs. lower quartile). A model including leg bioimpedance, age, sex, and self-reported history of myocardial infarction showed good predictive capacity of future heart failure hospitalization (C-index=0.82) and good calibration.

**Conclusions and Relevance:** Leg bioimpedance is inversely associated with heart failure incidence in the general population. A simple model of exclusively non-invasive measures, combining leg bioimpedance with history of myocardial infarction, age, and sex provides accurate predictive capacity.

**Key points:** *Question:* Which are the most important risk factors for incident heart failure?

*Findings:* In this population-based cohort study of ~500,000 individuals, machine learning identified well-established risk factors, but also several novel factors. Among the most important were leg bioimpedance and mean reticulocyte volume. There was a strong inverse relationship between leg bioimpedance and incident heart failure, also in adjusted analyses. A model entailing leg bioimpedance, age, sex, and self-reported history of myocardial infarction showed good predictive capacity of heart failure hospitalization and good calibration.

*Meaning:* Leg bioimpedance appears to be an important new factor associated with incident heart failure.

## Introduction

Heart failure is a common cause of mortality and is associated with substantial morbidity that greatly impacts quality of life of the affected individuals and their families. The lifetime risk of developing heart failure is high, and it is among the most common discharge diagnoses in the United States, with frequent costly readmissions for decompensation once disease has been established^1^. While invasively measured biomarkers have improved prognostication, prediction of new-onset heart failure remains challenging^2^. Increased knowledge of risk factors for incident heart failure in the general population could enable earlier diagnosis, preemptive treatment and risk factor control, to possibly mitigate disease development and improve outcomes.

We applied machine learning techniques in an agnostic search of new risk factors of incident heart failure in the UK Biobank, a population-based longitudinal cohort study of ~500,000 individuals, and found that leg bioimpedance was an important predictor of future heart failure. In a series of analyses, we further established that leg bioimpedance was independent of known risk factors for heart failure, and that this information is of high value for prediction of heart failure hospitalizations.

## Methods

### Study design and participants

Between 2006 and 2010, 502,639 individuals aged 49–69 years were enrolled in the UK Biobank, a longitudinal, population-based cohort study that recruited volunteers across 21 centers in England, Wales and Scotland. The study has been described in detail previously^3^, and online (https://www.ukbiobank.ac.uk). Briefly, participants have been carefully investigated, using a series of questionnaires, physiological measures, imaging, blood and urine biomarkers, and genotype data. Importantly, the UK biobank has been merged with the national in-hospital diagnosis and procedure registries, providing information about diagnoses prior to and after baseline of the study^3^.

For the main analyses, we excluded individuals with prevalent heart failure (either primary or secondary in-hospital diagnosis of heart failure, International Classification of Disease 10 [ICD10], I50; or self-reported history of heart failure, n = 2,188), leaving 500,451 individuals eligible. In secondary analyses of leg bioimpedance, we instead studied individuals with prevalent heart failure (n = 2,188). The UK Biobank study was approved by the North West Multi-Centre Research Ethics Committee and all participants provided written informed consent to participate in the UK Biobank study.

### Leg bioimpedance and other risk factors

In our phenome-wide scan of risk factors for incident heart failure, we considered all 3646 variables reflecting different aspects of lifestyle, health and disease-related factors in the UK Biobank (except a few administrative variables such as meta-data regarding the genomics part of UK Biobank, as well as the genomics data), as well as all ICD-10 diagnoses (excluding administrative, temporary, and unspecific codes [chapters U, V, W, X, Y and Z]) (**Supplementary Table 1**).

Leg bioimpedance was measured using the Tanita BC418MA body composition analyzer. In our confirmatory analyses, we defined leg bioimpedance as the mean of left and right leg bioimpedance.

Blood pressure and body mass index were measured in a standardized fashion; as were medical history, smoking status, alcohol consumption through standardized questionnaires (full details available at UK Biobank data showcase: http://biobank.ctsu.ox.ac.uk/crystal/). High alcohol intake was defined as “Daily or almost daily” intake of alcohol. Chronic kidney failure (N18) and coronary heart disease (I20–I25), were defined as presence of these (primary or secondary) ICD-10 diagnoses in hospitalizations prior to inclusion in UK Biobank.

### Outcome and follow-up

The primary outcome was hospitalization for heart failure (ICD-10: I50) as primary cause during follow-up. There were 1,054 incident events of heart failure during follow-up of up to 9.8 years, with a median follow-up of 6.2 years (lower and upper quartile, 5.5 and 6.9).

### Statistical analysis

For discovery of new potential risk factors for heart failure, we used a gradient boosting machine (GBM) model. This machine learning method allows the computer to learn how to classify individuals as having heart failure or not in an iterative fashion, considering all of the 3646 available variables, improving prediction by each iteration. In addition to providing a model for classification, it provides a measure of how important each variable is in this classification. We used hospitalization for heart failure (i.e. our primary outcome) as the response variable. The dataset was divided into training (60%), validation (20%), and test (20%) sets. The maximum tree depth was set to 3, the maximum number of trees to 50, and the learning rate to 0.01. Performance was assessed by the receiver operating characteristic area under the curve (ROC-AUC), and error was assessed by the mean squared error (MSE). Results are presented as variable importance for the classification.

Next, we assessed the relationships between the top 15 predictors of heart failure from the GBM model and incident heart failure by using Cox proportional hazards regression in two sets of models: a) adjusting for age and sex; and b) additionally adjusting for body mass index, systolic and diastolic blood pressure, anti-hypertensive treatment, alcohol consumption, diabetes, smoking status, prevalent chronic renal failure, and prevalent coronary heart disease. Continuous predictors were entered as restricted cubic splines to account for any non-linear associations. The association between predictors and heart failure are reported as hazard ratios (HR) and 95% confidence intervals (CI) for the upper quartile vs. the lower quartile for continuous variables, or for presence or absence for binary variables.

As leg bioimpedance was deemed of particular importance after our initial analyses, we proceeded with in-depth analyses of this variable. First, we performed a sensitivity analysis, where all individuals with any primary or secondary in-patient diagnosis in chapter nine of ICD-10 (Diseases of the circulatory system) were excluded, as well as anyone with any of the following self-reported conditions: hypertension, prior stroke or TIA, high cholesterol, diabetes mellitus, angina pectoris, or myocardial infarction (as well as self-reported heart failure) rendering a sub-sample of 309,079 individuals. Further, we assessed leg bioimpedance in the full population in relation to incident or prevalent heart failure, and to any incident or prevalent cardiovascular disease. Distributions are presented as density plots. The predicted rate of heart failure hospitalization up to 8 years after inclusion in the study was plotted in relation to leg bioimpedance in a spline plot. The x axis was truncated at the 0.1^th^ percentile at the lower end, and the 99.9^th^ percentile in the higher end. Normality was assessed with visual inspection of the distributions, and levels of leg bioimpedance were compared with t-tests.

The discriminatory ability of the models is presented using c-indices. We assessed calibration by visual inspection of calibration plots, and performed bootstrap validation using 100 bootstrap samples. All analyses were conducted using R versions 3.3.0 and 3.3.3, and the machine learning platform H2O version 3.10.4.1.

## Results

Baseline characteristics of the study sample are presented in **Table 1**. The 15 variables with highest importance in the GBM model are shown in **Figure 1**. Among these are some of the most well-established risk factors for heart failure, such as manifestations of coronary artery disease and type 2 diabetes; as well as known correlates of heart failure, such as obesity, microalbuminuria and anemia. There were also several novel predictors of heart failure. Among the most important were left and right leg bioimpedance and mean reticulocyte volume. The ROC-AUCs were similar in the training, validation, and test sets; 0.79, 0.79, and 0.81; as were the MSE; 0.0020, 0.0023, 0.0022, i.e. acceptable discrimination and low error, which were very similar in the three sets, suggesting that the model was not overfitted.

**Figure 1:**
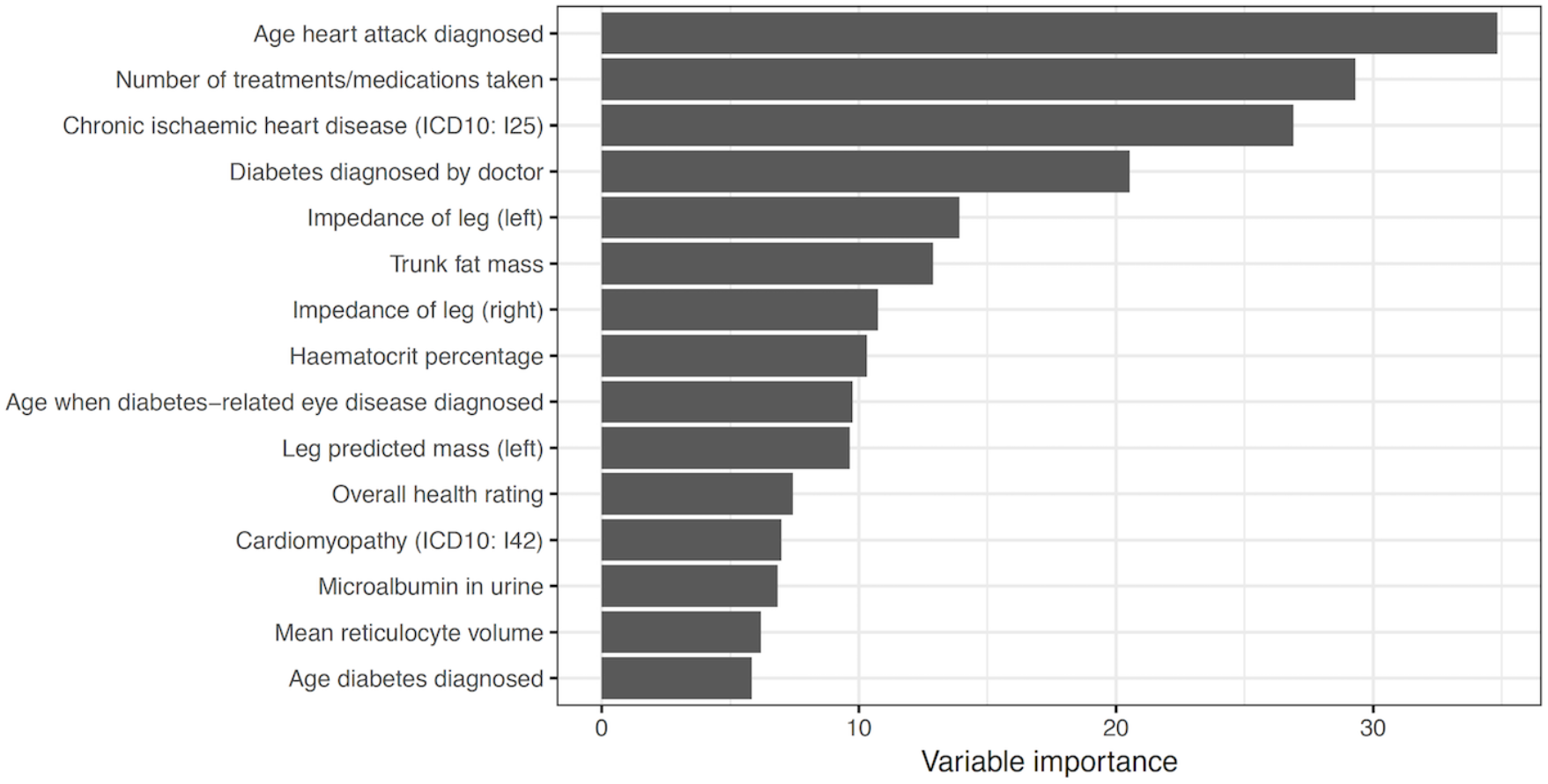
Variable importance for the top 15 variables in the gradient boosting machine model for incident hospitalization for heart failure.

**Table 1:** Baseline characteristics

| Characteristic | N |  |
| --- | --- | --- |
| Leg bioimpedance | 490,660 | 246.5 (223.5–270.5) |
| Age | 500,488 | 58 (50–63) |
| Male sex | 500,488 | 45% (227,533) |
| BMI | 497,382 | 26.7 (24.1–29.9) |
| Systolic blood pressure | 499,157 | 136.5 (124.5–149.5) |
| Diastolic blood pressure | 499,159 | 82.0 (75.3–89.0) |
| Antihypertensive medication | 492,594 | 21% (104,726) |
| Alcohol intake daily or almost daily | 498,969 | 21% (102,526) |
| Diabetes mellitus | 497,858 | 5% (25,862) |
| Current smoker | 497,576 | 11% (52,904) |
| Prevalent chronic renal failure | 500,448 | 0.1% (635) |
| Prevalent coronary heart disease | 500,448 | 4% (17,678) |

For continuous variables, medians and interquartile ranges are reported; for categorical variables, percentages and frequencies. N is the number of non-missing values.

For in-depth analyses of the most important variables, we then performed two sets of Cox regression models (age- and sex-adjusted; and adjusting for established heart failure risk factors) relating the top 15 predictors to incident heart failure during up to 9.8 years of followup (**Table 2**). Among the independently associated variables were well-established risk factors, such as ischemic heart disease and diabetes, but also other, novel markers, such as mean reticulocyte volume and leg bioimpedance.

**Table 2:** Cox models of the top 15 associated variables from the machine learning approach. “Diabetes diagnosed by doctor” and “Age diabetes diagnosed”, was instead analyzed as history of diabetes mellitus.

| Variable | Age- and sex-adjusted<br>HR (95%CI) | Fully adjusted**<br>HR (95%CI) |
| --- | --- | --- |
| Prior myocardial infarction* | 7.06 (6.06–8.23) | 4.29 (3.62–5.07) |
| Number of treatments/medications taken | 3.05 (2.44–3.82) | 1.96 (1.52–2.53) |
| Chronic ischemic heart disease (ICD10: I25) | 6.88 (5.95–7.95) | 4.04 (3.44–4.75) |
| History of diabetes* | 5.27 (4.60–6.03) | 2.45 (2.10–2.87) |
| Leg bioimpedance (left) | 0.59 (0.48–0.72) | 0.75 (0.60–0.94) |
| Trunk fat mass | 1.73 (1.43–2.09) | 0.86 (0.64–1.17) |
| Leg bioimpedance (right) | 0.60 (0.49–0.73) | 0.78 (0.62–0.97) |
| Hematocrit (%) | 0.56 (0.47–0.67) | 0.73 (0.61–0.87) |
| Diabetes-related eye disease* | 8.57 (6.01–12.21) | 2.01 (1.37–2.94) |
| Leg predicted mass (left) | 3.73 (2.74–5.07) | 1.45 (0.99–2.12) |
| Overall health rating | 3.14 (2.91–3.38) | 2.04 (1.86–2.22) |
| Cardiomyopathy (ICD10: I42) | 18.73 (11.59–30.27) | 9.61 (5.70–16.18) |
| Microalbumin in urine (log) | 2.12 (1.62–2.77) | 1.83 (1.38–2.42) |
| Mean reticulocyte volume | 1.28 (1.10–1.50) | 1.19 (1.01–1.39) |

\* Age when occurrence of myocardial infarction, diabetes, and diabetes-related eye disease were diagnosed were dichotomized to reflect the presence of these conditions or not.
\*\* Fully adjusted model includes age, sex, BMI, systolic blood pressure, diastolic blood pressure, antihypertensive treatment, alcohol consumption, diabetes, smoking status, prevalent chronic renal failure, prevalent coronary heart disease.
CI = Confidence interval
HR = Hazard ratio. For continuous variables, HR represents upper vs. lower quartile.
ICD = International classification of disease

As left and right leg bioimpedance emerged as novel, independent risk factors and strong predictors of incident heart failure that can be assessed in a simple and non-invasive manner, we decided to focus remaining analyses on leg bioimpedance, defined as the mean of left and right leg bioimpedance. The distribution of leg bioimpedance in relation to incident heart failure was investigated, and it showed fairly good separation between those who experienced future outcome compared with those who did not (**Figure 2**; p = 1.1 x 10^-72^). Compared with individuals who were free from heart failure during follow-up, individuals with incident heart failure had significantly lower leg bioimpedance (**Figure 3**).

**Figure 2:**
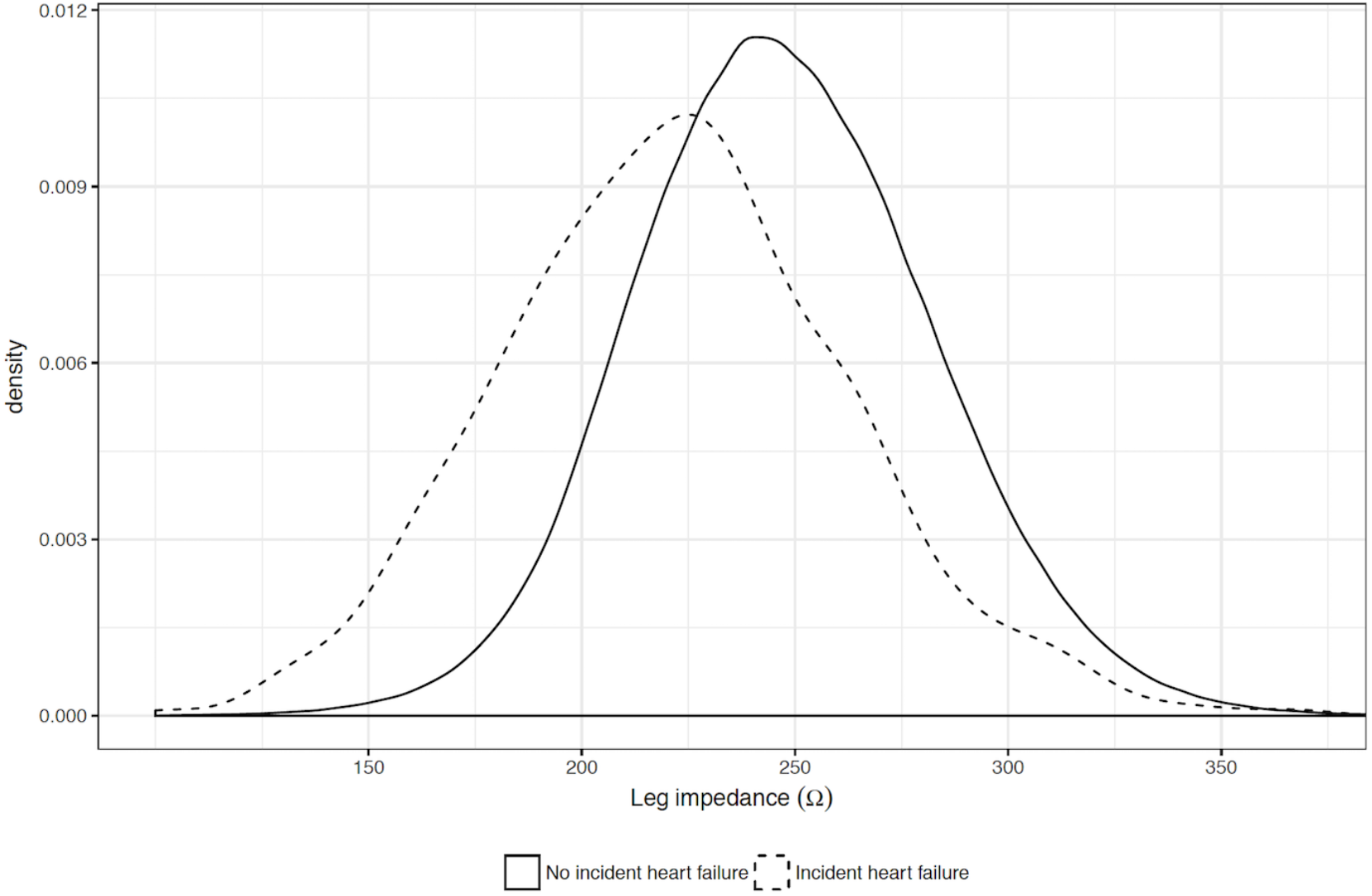
Distribution of leg bioimpedance in relation to incident heart failure.

**Figure 3:**
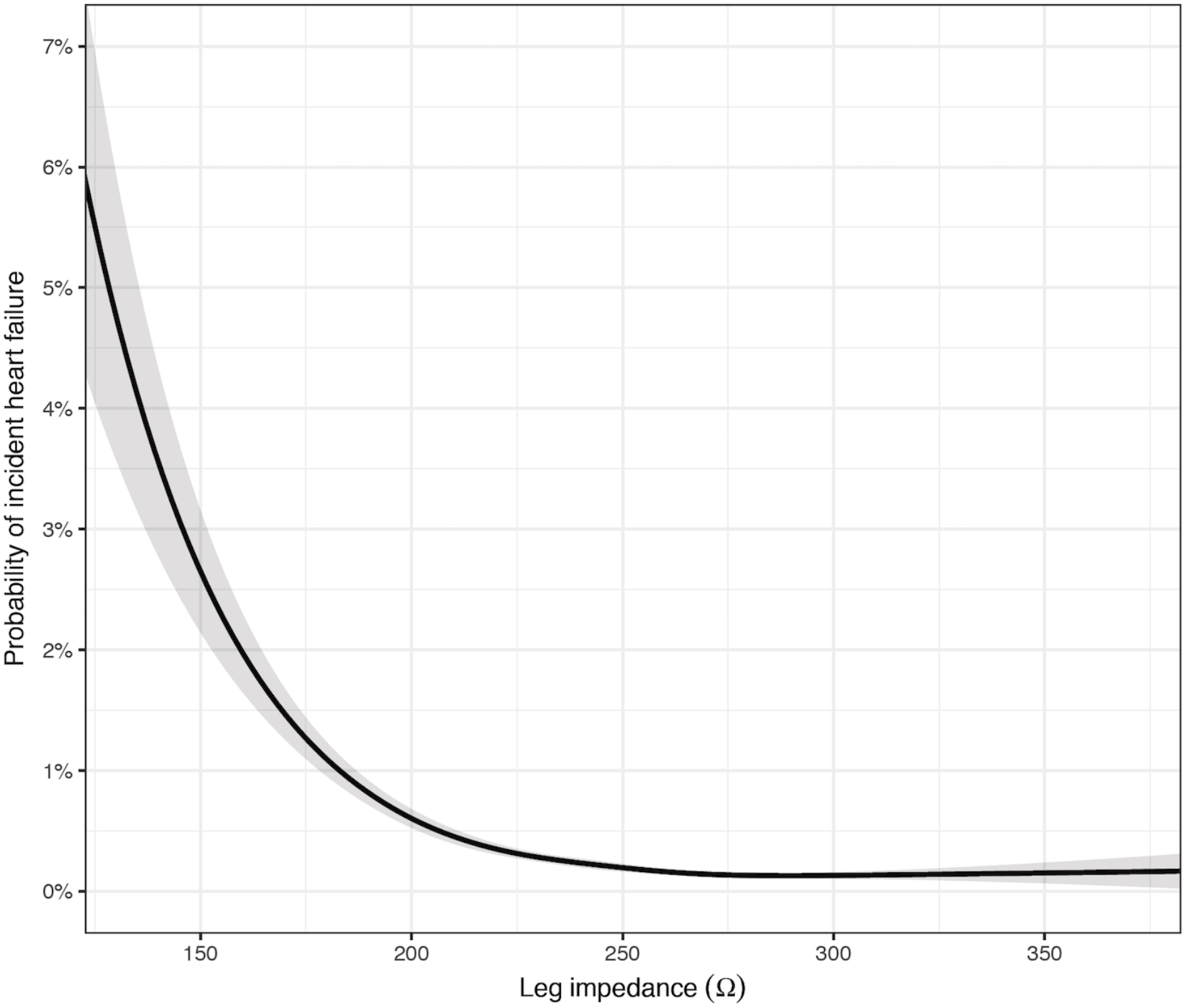
Spline plot of probability for heart failure hospitalization in relation to leg bioimpedance.

In sensitivity analyses, we assessed leg bioimpedance in relation to incident heart failure in a subsample, excluding all individuals who had prior hospitalizations for any cardiovascular disease, as well as those with self-reported presence of cardiovascular disease and/or cardiovascular risk factors (N=309,079 with 212 cases [0.07%] of incident heart failure). In this healthy subsample, associations of leg bioimpedance with heart failure were non significant, but displayed similar effect sizes as in the full study sample (hazard ratio, 0.74; 95% confidence interval, 0.47–1.15; comparing upper vs lower quartile in a multivariable-adjusted model; **Figure S1**).

In additional sensitivity analyses in relation to prevalent cardiovascular disease, leg bioimpedance was consistently lower in those experiencing heart failure hospitalizations during follow up (**Figures S2–S3** in Supplement), and in those with known prior heart failure, lower leg bioimpedance was associated with having multiple re-hospitalizations during follow-up (**Figure S4** in Supplement).

When adding leg bioimpedance to a model including only age and sex, the C-index increased from 0.76 to 0.80, which should be compared with an increase from 0.76 to 0.79 when adding self-reported myocardial infarction to the model (**Table 3**). Combining age, sex, leg bioimpedance and self-reported myocardial infarction in a model led to a C-index of 0.82. This model was well-calibrated (calibration slope, 0.9985; **Figure S5** in Supplement), and the optimism-corrected C index was 0.82 (i.e. essentially unchanged, obtained via bootstrap validation), indicating that overfitting is not an issue. When taking known risk factors for heart failure into account, leg bioimpedance remained significantly inversely associated with incident heart failure, calibration was similar as for the smaller model (**Figure S6** in Supplement), and the C-index improved further to 0.85 (**Table 3**).

**Table 3:** Predictive capability of multivariable models of incident heart failure during up to eight years of follow-up.

| **Model** | **HR (95%CI)**<br>**Upper vs. lower**<br>**quartile of leg**<br>**bioimpedance** | **C-index** |
| --- | --- | --- |
| Age + sex | - | 0.76 |
| Age + sex + leg bioimpedance | 0.60 (0.48–0.73) | 0.80 |
| Age + sex + self-reported MI | - | 0.79 |
| Age + sex + self-reported MI + leg bioimpedance | 0.60 (0.49–0.74) | 0.82 |
| Fully adjusted* | 0.75 (0.59–0.94) | 0.85 |

\* Fully adjusted model includes age, sex, BMI, systolic blood pressure, diastolic blood pressure, antihypertensive treatment, alcohol consumption, diabetes, smoking status, prevalent chronic renal failure, prevalent coronary heart disease.
C-index = Concordance index, CI = Confidence interval, HR = Hazard ratio, MI = myocardial infarction.

## Discussion

Using a contemporary, prospective cohort study of >500,000 individuals from the general population, we have performed an extensive investigation of factors associated with incident heart failure. Our key findings are several-fold. First, using an agnostic machine learning approach in a phenome-wide manner, we established the strongest predictors of incident heart failure, which included well-established (e.g. coronary artery disease, type 2 diabetes), previously suggested (e.g. obesity, microalbuminuria, anemia) and novel (e.g. leg bioimpedance, mean reticulocyte volume) potential risk factors of heart failure. Second, we report that leg bioimpedance was inversely associated with incident heart failure, even after taking known risk factors into account in multivariable-adjusted Cox regression models. Third, a multivariable model including leg bioimpedance, age, sex and self-reported prior myocardial infarction provided accurate prediction of heart failure incidence, highlighting the potential of this simple, non-invasive, and cheap measure for clinical use.

### Previous Studies of Bioimpedance and Heart Failure

To our knowledge, this is the largest study of risk factors for heart failure hospitalization, and the only study of bioimpedance in relation to incident heart failure in the general population. There are a few previous studies that have assessed bioimpedance in relation to heart failure decompensation in patients with already established severe heart failure treated with cardiac resynchronization devices or implantable defibrillators, where impedance could be measured using the pacemaker leads. In that setting, decompensation is characterized by a decrease in intrathoracic impedance, indicative of volume overload, well before re-hospitalization for congestion^4^. There are also a few prior studies of non-invasive impedance measures in patients with established heart failure. In a phase II trial, treatment guided by lung impedance (assessed using external electrodes) reduced heart failure hospitalizations^5^. In addition, bioimpedance measures have been suggested as an adjunctive test for patients presenting with dyspnea at the emergency department, to identify heart failure patients when other test are equivocal^6,7^. Whether the information obtained from impedance measures in patients with established heart failure translates into improved outcomes is, however, uncertain^5,8,9^.

### Strengths and Limitations

Strengths of our study include the unprecedented sample size from the general population with consistent assessment of risk factors for heart failure, including leg bioimpedance in >500,000 individuals; the minimal loss-to-follow-up; and our stringent, yet exploratory approach that allowed us to discover leg bioimpedance as a novel risk factor for heart failure in the general population. Our study also has some important limitations. Cardiac-specific biomarkers, especially the natriuretic peptides which are strongly associated with incident heart failure when screening the general population^10,11^, are not available in the UK Biobank. The factors included in our prediction models do not require any blood draw, and could be easily and rapidly assessed in a primary care facility using simple and available instruments and self-reported information. However, it would have been interesting both from a mechanistic and predictive standpoint to be able to study leg bioimpedance in relation to, for example B-type natriuretic peptide. Furthermore, incident heart failure was defined as a diagnosis of heart failure in national registries during follow-up, rather than via review of journal records including echocardiography reports. As a consequence, the etiology of the heart failure, and whether the events represented heart failure with reduced or preserved ejection fraction could not be discerned, and milder heart failure never requiring hospitalization may not have been captured. Finally, even if the association between leg bioimpedance and incident heart failure was strong also when excluding individuals with any prior cardiovascular diagnoses or self-reported cardiovascular history and risk factors (including self-reported heart failure, which presumably would capture also milder cases of heart failure), reverse causation is always a possibility in any observational analysis. In this case, we cannot be certain whether the associations are attributable to subclinical heart failure, as impedance changes have been suggested to precede heart failure symptoms^12^, or if this relates to specific biological attributes associated with increased risk of future heart failure. Along the same lines, even if we have adjusted our models for known risk factors for heart failure, there is certainly risk of residual confounding, so any conclusions regarding mechanistic and causal relations between leg bioimpedance and heart failure needs to be studied using other study designs. That said, our findings that leg bioimpedance is a strong predictor of heart failure with good discrimination and calibration in the general population remain valid regardless of the above limitations.

## Conclusions

Leg bioimpedance is inversely associated with heart failure incidence in the general population. A model combining leg bioimpedance with self-reported history of myocardial infarction, age, and sex shows accurate predictive capacity that may be useful in the clinical setting, provided independent prospective validation.

## Acknowledgements

This research has been conducted using the UK Biobank Resource (^3^, and online (www.ukbiobank.ac.uk) under Application Number 13721. Computing for this project was performed on the Sherlock cluster. We would like to thank the Stanford Research Computing Center for providing computational resources and support that have contributed to these research results.

## Author Contributions

Study concept and design: Lindholm, Leeper, Ingelsson Acquisition, analysis, or interpretation of data: All authors Drafting of the manuscript: Lindholm, Ingelsson Critical revision of the manuscript for important intellectual content: All authors Statistical analysis: Lindholm, Ingelsson Study supervision: Leeper, Ingelsson

